# PathPinpointR: Predicting the progression of sc-RNAseq samples through reference trajectories

**DOI:** 10.64898/2026.04.21.715327

**Authors:** Moi T. Nicholas, Disha Mehta, John. F. Ouyang, Ahmed Dawoud, Charlotte Ellison, Juri Westendorf, Luke Green, Paul Skipp, Owen J. L. Rackham

## Abstract

Single-cell RNA sequencing (scRNA-seq) has transformed our ability to analyse cellular heterogeneity, enabling detailed mapping of cellular progression. Trajectory inference tools construct trajectories from scRNA-seq data, facilitating the tracing of cellular progression through developmental pathways. PathPinpointR (PPR) is a lightweight and user-friendly R package developed to predict and compare the positions of scRNA-seq samples along reference biological trajectories, such as those created from large cell atlas projects. PPR utilises sets of switching-gene events from reference trajectories as indicators of cellular progression. By applying these positional indicators to query datasets, each cell can be accurately assigned a pseudo-time value, providing predictive insight into its position along a trajectory. This information can be used to stage cells within an established developmental process, or to evaluate how different patient samples compare when mapped onto reference disease or drug response trajectories.

**Availability:** PathPinpointR is available at https://github.com/moi-taiga/PathPinpointR.

## 1. Introduction

Single-cell RNA sequencing (scRNA-seq) has become essential for exploring cellular heterogeneity. It provides a high-resolution view of processes such as cell development or disease progression. Recent international efforts to collect large cell atlases (Regev *et al*., 2017) are uncovering novel cell types (Warner van Dijk *et al*., 2024), as well as providing new ways to investigate (Rood *et* al., 2025) and diagnose disease (He *et al*., 2021). However, as these atlases grow in size and complexity, ways to leverage them to increase the value, insight and power of smaller studies are needed. Here, we present PathPinpointR, a tool that leverages cell atlas–based reference trajectories to quickly and efficiently contextualise the data generated in smaller studies.

Trajectory inference tools trace cells’ progression through biological pathways by assigning cells a pseudotime value – a metric of cellular order according to progression along a given pathway (or process). These pathways, defined by pseudotime, are referred to as trajectories. Trajectories represent a path in gene-expression space along which cells progress, defined by gene expression. They are used to capture processes such as cell differentiation, disease advancement, and in vitro perturbation effects (Liu *et al*., 2020; Paik *et al*., 2020).

The changes in gene expression which drive or are caused by progression along a trajectory can be identified with tools such as Switchde and GeneSwitches (GS) (Campbell and Yau, 2017; Cao *et al*., 2020). These tools highlight switching-genes, defined as genes that transition from one expression state to another (e.g., from on to off) over the course of a trajectory. Individually, these switching-gene events serve as markers of cellular progression; collectively, the entire set (which we refer to as the switching-gene set) can be used to define the trajectory.

Much like the sets of cell-type-specific marker genes which are now routinely used to classify distinct cell types in single-cell RNA sequencing studies, extracting switching-gene sets from large reference datasets in such a way that they can be used to better characterise smaller or patient-specific datasets represents a significant opportunity.

To enable this, we developed PathPinpointR (PPR) to utilise trajectory-specific switching-gene sets from reference datasets, as a way to predict the pseudotime of individual cells within query datasets. Existing approaches for achieving this typically require integrating query data with the reference data and reapplying trajectory inference algorithms; PPR uses the switching-gene events as checkpoints to rapidly predict a pseudotime for each cell within a query dataset. By aggregating these cell-level predictions, PPR delivers predictive insights into each query sample’s overall position along the trajectory. Furthermore, PPR can compare predictions from multiple query datasets across different experimental conditions, providing a way to order samples based on their relative progression through a trajectory.

Taken together, PPR combines efficiency with ease of use, enabling researchers to rapidly and easily estimate and compare the pseudotime of cells across diverse biological trajectories. We demonstrate the utility of PPR by applying it to simulated data and a developmental time-series of blastocyst formation.

## 2 Implementation

### Data Preprocessing

When applying PPR, two inputs are required: (i) scRNA-seq (gene-cell matrix) from a sample of interest (query data) (ii) a reference trajectory (reference data). Key to the successful prediction of sample position is selecting a reference dataset with sufficient cells to represent the chosen biological pathway accurately. The reference dataset should capture the diversity of cellular states along the trajectory.

Both the query and reference data must undergo some preprocessing steps before being used with PPR. Firstly, a trajectory inference method must be applied to generate pseudo-time values for each cell in the reference dataset, producing an ordered sequence that reflects cellular progression. Secondly, the gene expression values from both the reference and query datasets must be binarised (0/1). Finally, the switching genes present in the reference trajectory must be identified, together with a measure of their confidence as switching genes, in order to create the trajectory-specific switching-gene set.

### Pseudotime prediction of cells within query data

The pseudotime prediction of the query data happens first at the cell level, where the position of each cell is predicted. This is achieved by considering the expression, or lack thereof, for each gene in the trajectory-specific switching-gene set. For each gene in the set, the expression status of an individual cell is processed according to the logic represented in Figure 1. In summary, if a gene is expressed in a given cell and this gene is known to activate during the trajectory, all positions beyond that pseudotime are considered plausible. Conversely, if the gene is not expressed, all positions prior to the switching pseudotime are considered plausible.

**Figure 1.**
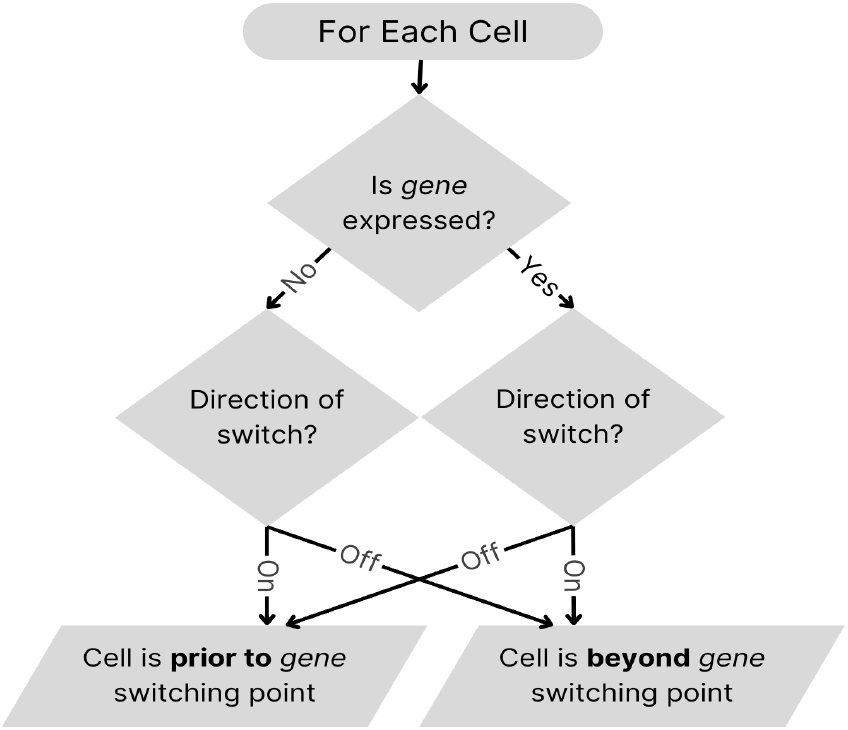
Flowchart depicting the logic underpinning PathPinpointR’s positional prediction. This cell-level decision process is evaluated for each switching gene across every cell in a query dataset. The algorithm first assesses whether a target gene is expressed, then identifies if that gene switches on or off along the trajectory, and finally determines whether the cell is positioned before or after the gene’s switching point.

For every gene in the switching-gene set, a binary variable is set for each pseudotime point (plausible/not plausible). By summing this across the entire set of switching genes, an estimate of the most plausible pseudotime for that specific cell can be generated.

### Performance Metrics

To estimate accuracy for a given trajectory-specific switching-gene set, PPR compares the predicted position of cells from the reference data to the pseudo-time produced by a trajectory inference (see function `accuracy_test()` for full details). The difference per cell is considered the inaccuracy, with the mean inaccuracy representing a global performance measure that can be minimised (see Figure 2).

**Figure 2.**
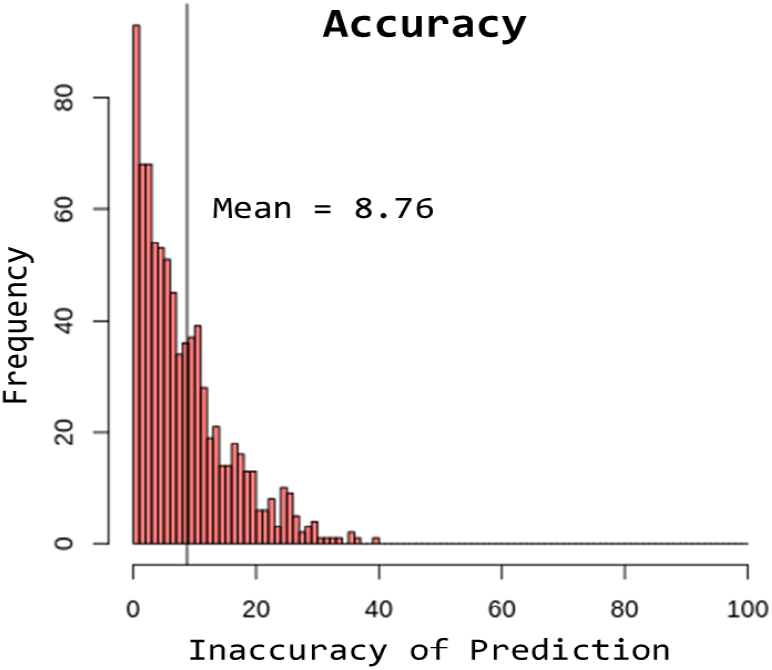
Graphical output of accuracy_test() Histogram of the inaccuracy of predictions when running PPR on cells from the reference trajectory. Inaccuracy, being the difference between the predicted pseudotime and the true pseudotime of cells from the reference; the mean inaccuracy is highlighted as 8.76.

To identify the optimum size of a trajectory-specific switching-gene set, PPR reports the mean inaccuracy from runs using increasing numbers of switching genes. By default, the set size that provides the lowest mean inaccuracy is reported, as in Figure 3.

**Figure 3.**
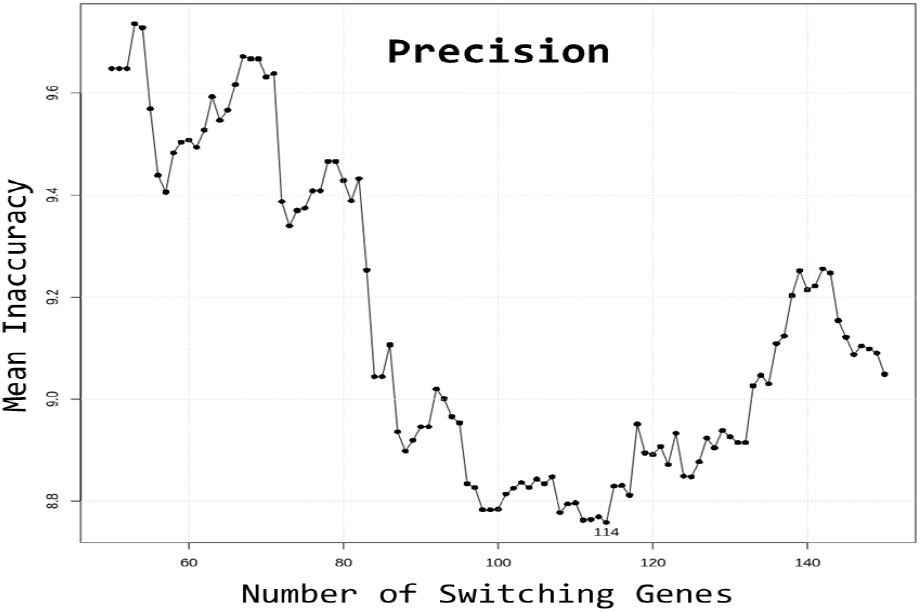
Graphical output of precision() Average inaccuracy by the number of switching genes used. 114 is found to produce the minimum inaccuracy; this value would be selected for further PPR analysis for the given trajectory.

### Aggregate pseudotime prediction for query data

To generate a sample-level prediction, PPR aggregates positional indicators from individual cells to estimate the query’s most likely position along a trajectory. Summing the plausibility indicators for each gene across all cells yields a vector representing the frequency of these gene-level predictions along pseudotime. This distribution can be visualised as a histogram, as shown in Supplementary Figure S1.

## 3 Vignettes

To validate PPR, it was tested both on subsets of reference data and on true sample data. Two examples are given: one using synthetically generated data, and one measuring early development in a blastocyst sample. Both are available via GitHub.

### Synthetic data

Synthetic data was generated using a method built upon that demonstrated by the Slingshot vignette (Street *et al*. 2024). Simulated expression values were created for 1,255 genes across 300 cells, organised into groups with distinct expression patterns. These groups included non-differentially expressed genes and differentially expressed genes, which switch on or off at various points along the trajectory. The methods used to generate the synthetic expression data for each group are described further in the supplementary data.

Two proxy samples were produced by taking subsets of the synthetic reference data, from early and late in the trajectory. PPR was accurate when predicting pseudotimes for cells from the reference data, and correctly identified the position of the two proxy samples. This serves as a proof of concept for PPR.

### Blastocyst development validation

The reference dataset used was comprised of multiple studies representing blastocyst development, integrated into a single Seurat object (Tyser *et al*. 2021; Sozen *et al*. 2021; Molè *et al*. 2021; Yanagida *et al*. 2021; Xiang *et al*. 2020; Zhou *et al*. 2019; Petropoulos *et al*. 2016). The reference data was sampled across multiple days, varying from Day 5 to Day 19, and subset to produce a trajectory of epiblast cell development. Two more blastocyst datasets were subset to only include epiblast cells and used as input samples for analysis. PPR was used to predict the stage of development for both test set blastocyst samples. Both samples were correctly identified as being from days 6 and 7, as observed in Figure 4.

**Figure 4.**
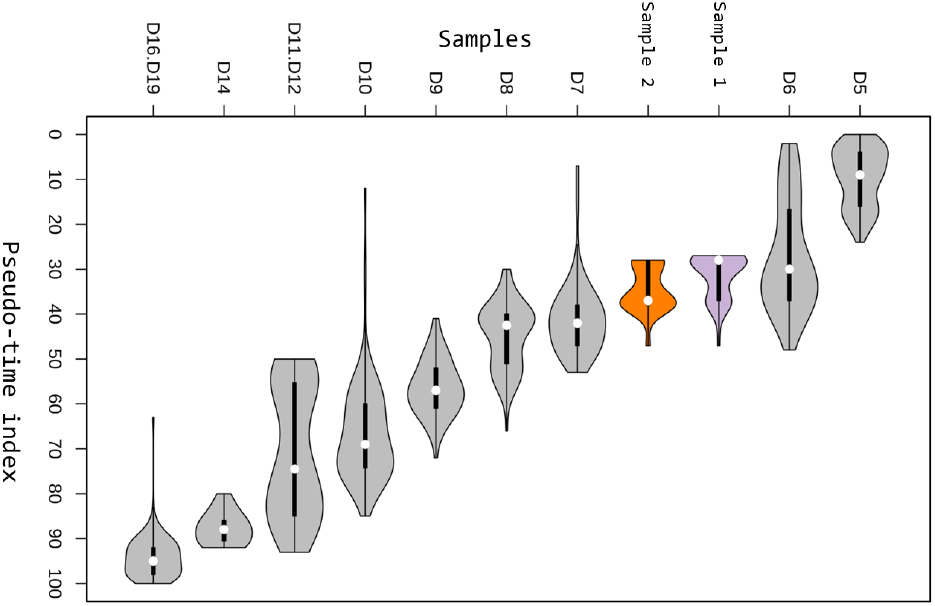
PathPinpointR prediction of epiblast pseudo-times. Horizontal violin plots representing the predicted position of two epiblast samples in the context of a development trajectory. Grey represents the true pseudotime of cells in the reference, split by day of development (D5, D6, …). Purple and Orange represent the predicted pseudo-time for cells in samples 1 and 2, respectively. The two samples are shown to be predicted to be at days 6 & 7 of development.

## Acknowledgements

The authors acknowledge the use of the IRIDIS High Performance Computing Facility and associated support services at the University of Southampton in the completion of this work.

We acknowledge the assistance of ChatGPT (OpenAI) and Gemini (Google) for discussions and writing support in developing sections of this manuscript.

