## Supplementat data for "PathPinpointR: Predicting the progression of sc-RNAseq samples through reference trajectories"

### Index:

Input data:

- Preparing a reference Trajectory
- Preparing Sample Data
- Blastocyst Vignette
- Slingshot Vignette

PPR Functions:

- predict_position
- Plotting with PPR
- accuracy
- precision

Slingshot Vignette:

- https://github.com/moi-taiga/PathPinpointR/blob/master/slingshot.md

Blastocyst Vignette:

- https://github.com/moi-taiga/PathPinpointR/blob/master/README.md

### Input Data:

#### Preparing a Reference Trajectory:

*Data Selection:*

A reference trajectory represents the biological process of interest, such as cell differentiation or disease progression. Firstly, a suitable dataset must be selected; the dataset must accurately represent the biological process of interest. Sufficient cells must be present to capture the diversity of cellular states and the relevant cell types or stages for the biological question being addressed.

*Trajectory Inference:*

Following data selection, a trajectory inference tool must be applied to provide cells with a pseudotime value. The choice of TI tool should depend on the dataset being analysed, Slingshot (Street et al. 2018) integrates particularly well with GeneSwitches (Cao et al., 2020).

*GeneSwitches:*

Next, the switching genes must be identified. These are genes that transition between binary expression states, as cells progress through the trajectory. To achieve this, GeneSwitches is used.
The first step is binarising gene expression, where expression values are converted to values of 1 for expressed genes and 0 for non-expressed genes. From the binarised expression matrix, switching genes can then be identified using the function `GeneSwitches::find_switch_logistic_fastglm`.

*Precision:*

To refine the list of switching genes, `PPR::precision` is applied. This function determines the optimal number of switching genes for analysis by evaluating their impact on prediction accuracy.

*Filter:*

Once the optimal number of switching genes is identified, `GeneSwitches::filter_switchgenes` is used to output a data frame containing the selected switching genes and their switching positions along the trajectory. The `topnum` argument should be set to the number of switching genes identified as optimal by `PPR::precision`, and the `r2cutoff` should be set to 0 to avoid excess filtering.

#### Preparing a Sample Dataset:

The sample dataset is the scRNA-seq data to be positioned on the reference trajectory. Preparation of the sample data beyond normalisation and filtering is minimal, the gene expression matrix must be binarised using GeneSwitches.

### Input data preparation for README.md:

The reference trajectory for this vignette was produced by integrating data across 7 studies (Tyser et al. 2021; Sozen et al. 2021; Molè et al. 2021; Yanagida et al. 2021; Xiang et al. 2020; Zhou et al. 2019; Petropoulos et al. 2016).

The sample data was extracted from PMID: 37683605 and was filtered to only include epiblast cells.

### Input data preparation for slingshot.md:

The function `PPR::get_synthetic_data()` generates a simulated single-cell gene expression dataset designed for use in the `slingshot.md` vignette. This dataset models a trajectory of cells ordered by pseudotime, following the method outlined in the Slingshot tutorial (Street et al. 2024). The purpose is to create an expression matrix that reflects distinct patterns of gene activity to mimic those observed in biological processes.

The generated expression matrix consists of genes (rows) and cells (columns) ordered by pseudotime. The genes are grouped by expression pattern, four gene expression patterns are simulated:

- **Baseline genes:** These genes exhibit minor variations in expression but do not undergo distinct activation or deactivation events. Their expression values remain relatively constant, and are generated by repeating fixed values to ensure little variation across pseudotime.
- **Switching genes:** These genes either activate or deactivate at specific points along the trajectory. The following equation represents the method used to determine expression values:

$$e^{arctan\left(k-\frac{N}{x}\right)}$$

  $N$ is the total number of cells (300).
  $k$ is a vector of integers ranging from $N$ to 1 (for deactivation) or 1 to $N$ (for activation), controlling the direction of change.
  $x$ determines the pseudotime at which the gene switches. For example, if $x$=2, the gene switches at the midpoint of the trajectory.

  The arctan function smooths the transition in gene expression in order to produce gradual -more biologically plausible- change. The exponential (e) is used to ensure all expression values are positive, as well as introducing a smoother and more pronounced change.

- **Transient genes:** These genes activate for a short duration before returning to an inactive state. Their expression is simulated in a three-phase structure: an increase of expression, a sustained peak, and a decrease in expression. The method of generating the expression of the increase and decrease is similar to that of the switching genes. The sustained peak is generated by a repeated value of 3, similar to the baseline genes.

These gene sets ensure that the dataset captures a range of gene regulation scenarios commonly observed in single-cell RNA-seq data. Random variation is introduced to the gene expression matrix in order to produce variation between genes within a set.

The simulated count data is organised into a `SingleCellExperiment` object, with rarely expressed genes removed and raw counts normalised. Principal Component Analysis (PCA) is applied, and each cell is assigned to one of six numerically labelled clusters. The function `PPR::get_synthetic_data` returns the `SingleCellExperiment` object containing the simulated count data, normalised values, reduced dimensions (PCA), and cluster assignments.

### PPR Functions:

#### Predict_position:

The `predict_position` function uses the expression level of switching genes in individual cells to estimate their positions along a biological trajectory. It then aggregates these cell-level positions to provide an overall position for the sample.

The function requires two inputs. First, a sample provided as a SingleCellExperiment object, which must contain a binarised gene expression matrix. Second, a data frame that details the switching genes; the data frame must indicate the pseudotime at which each gene is expected to switch, and whether the gene is turning “up” or “down”.

For each cell, the function creates a matrix with rows corresponding to genes and columns representing 100 pseudotime indices. It uses each gene’s switching time and direction of switch to populate the empty matrix with probable pseudotime positions (indicated by a value of 1). For example, if a switching gene switches “up” during a trajectory and is expressed in the cell, the function marks all positions (columns) from the switching time onwards as 1. If –for the same gene– a cell had no expression, then all pseudotime indices prior to the switching time would be marked as 1. The inverse is true for switching genes with a “down” direction.

Once each cell’s gene-by-pseudotime matrix is populated, these matrices (termed genomic_expression_traces) are summarised. First, each cell’s matrix is reduced to a single row of column sums –providing an overall likelihood of the cell being at each pseudotime point– they are then combined into one matrix (cells_flat). Then, these cell-level summaries are aggregated further into a single vector (sample_flat), which represents the sample’s overall trajectory position.

The function returns a PPR object, a structured list containing genomic expression traces for individual cells, summarised trajectory probabilities for each cell, and an overall trajectory summary for the sample.

#### Plotting with PPR:

The plotting functionality of PPR is broken down into smaller functions:

The `ppr_plot` function is the foundational plotting function of PPR, designed to create a base plot for visualising pseudotime and PPR scores. It takes no inputs and outputs an empty plot with a predefined range for both the x and y axes. The x-axis is labelled as "Pseudo-Time Index", and the y-axis is labelled as "PPR Score". This plot serves as a canvas upon which additional visual elements can be overlaid using other functions.

The `sample_prediction` function is used to plot the predicted pseudotime position of a single sample within the plot created using `ppr_plot` . This function takes as input a `PPR_OBJECT`, which contains the PPR data for the sample. Additionally, it accepts optional parameters for setting the colour of the line and annotations (default is "red"), as well as a label string for the sample (default is "sample name"). The output of this function overlays a line showing the PPR scores across pseudotime, with a dashed line indicating the predicted pseudotime position. A label is added at the point where the sample has its maximum PPR score, offering further clarity on its location within the pseudotime trajectory. It can be run multiple times to overlay multiple samples on one plot.

The `reference_idents` function overlays a boxplot showing the pseudotime distributions of reference cells, grouped by their identity. The function takes as input a `SingleCellExperiment` object (`reference_sce`), which represents the reference trajectory, and a column name from the `colData` of `reference_sce` that specifies the cell groupings. The output of this function is a series of horizontal boxplots at the top of the PPR plot, illustrating the pseudotime distributions of reference cells by group. Labels for each group are displayed on the y-axis for easy identification.

The `switching_times` function highlights pseudotime points where genes switch on or off. It requires a character vector of gene names as input, along with the `switching_genes` output, which is derived from the `GeneSwitches::find_switch_logistic_fastglm()` function. This output contains the switching pseudotime indices for the genes of interest. The function adds dashed lines at these pseudotime switching points and labels the lines with the corresponding gene names, enabling the user to visually track the transitions of specific genes over pseudotime.

The `cell_position_box` function creates a boxplot overlay that visualises the pseudotime positions of cells within a sample. It requires a `PPR_OBJECT` containing the PPR data for the sample as input, along with a colour specification for the boxplot. The output of this function is a horizontal boxplot visualising the pseudotime positions of cells, with the option to show individual cell positions as points, allowing for a more granular view of cell distribution across pseudotime.

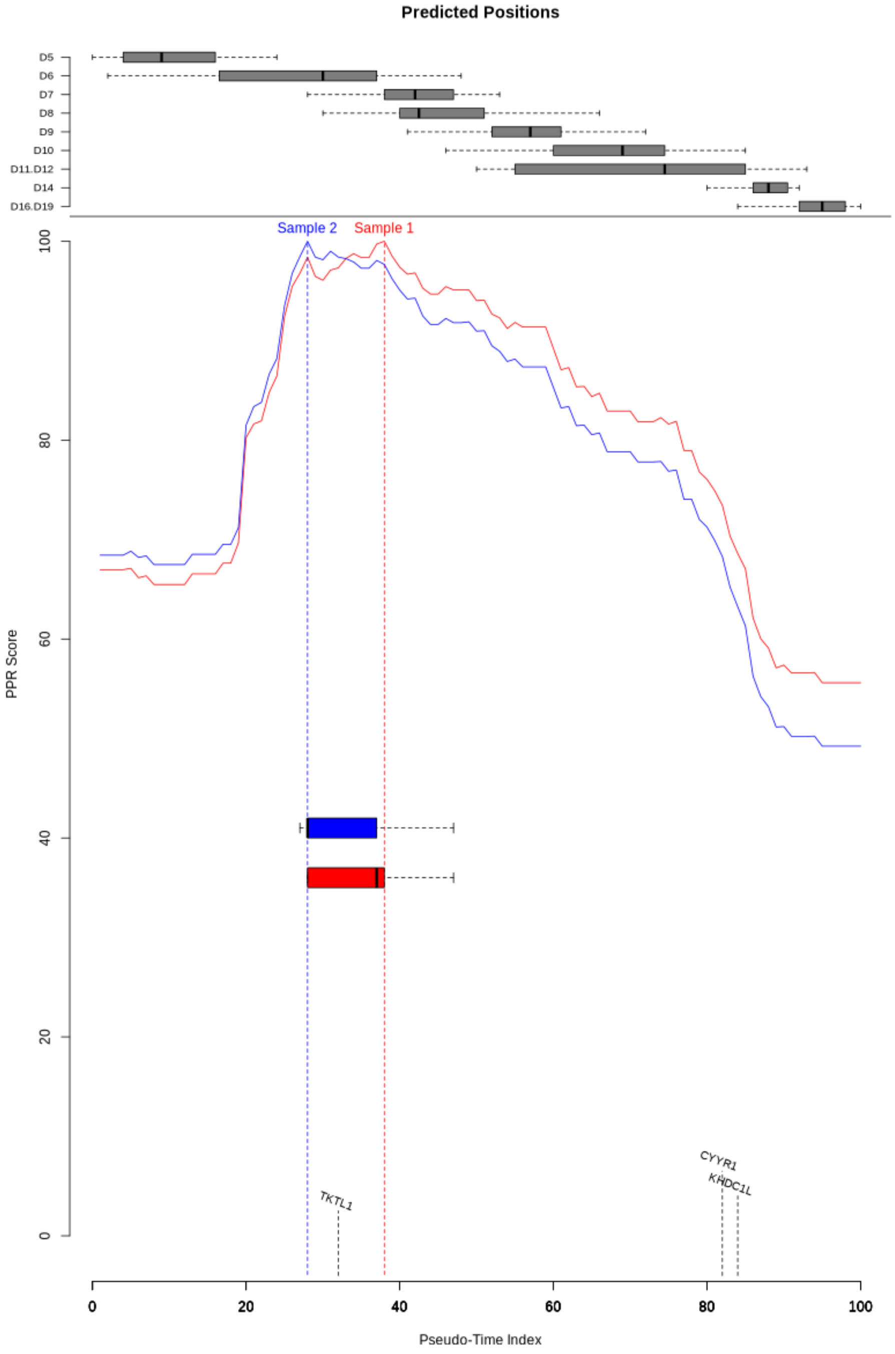


**Figure S1.** Visualisation of pseudotime predictions in PathPinpointR.
The upper section displays boxplots of reference cell distributions across pseudotime, grouped by identity (in this case days of blastocyst development). The lower section shows PPR scores distributed across pseudotime for two samples (red and blue), with vertical dashed lines marking their predicted pseudotime positions. The Y-axis represents the PPR score, derived from the sum of genes positively influencing the predicted position. Using the switching_times function, three switching genes have been labelled on the x-axis.

### Accuracy:

The `accuracy_test` function evaluates how well PPR predicts the positions of reference cells in pseudotime by comparing its predictions to the known pseudotime values from reference data.

It takes as input a `PPR_OBJECT` (which contains predicted cell positions) and a `SingleCellExperiment` object (`reference_sce`) that holds the true pseudotime values for each cell. The function then calculates the difference between the predicted and true positions, providing a measure of accuracy.
If `plot = TRUE`, the function generates a histogram showing the distribution of inaccuracies, highlighting how far predictions deviate from the actual values.
If `random = TRUE`, the function also computes random predictions and their inaccuracies, offering a baseline comparison.
When `plot = FALSE`, the function returns a data frame containing the results.

In summary, `accuracy_test` provides both numerical and visual assessments of prediction accuracy, allowing the user to quantify the efficacy of PPR when using a given reference data set.

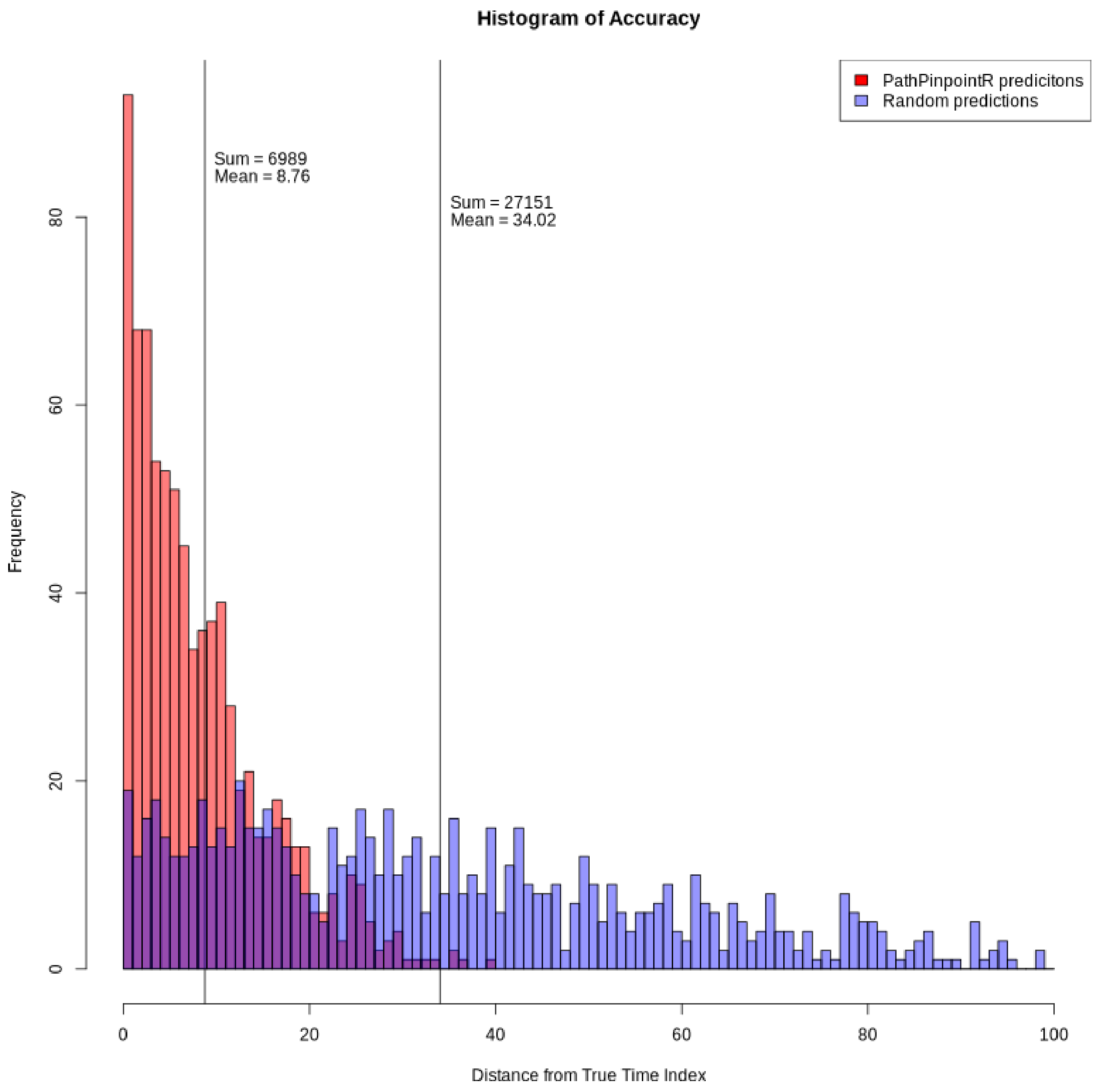


**Figure S2.** Distribution of prediction inaccuracies in PPR.
The histogram displays the distribution of inaccuracies, the distance in pseudotime from a cell's predicted position to its "true" pseudotime as assigned by a trajectory inference tool. PPR predictions are shown in red, and random predictions (if `random = TRUE`) are shown in blue. The sum of inaccuracies and the mean for each group is shown at their respective means.

### Precision:

The `precision` function is designed to determine the optimal number of switching genes for use in the PPR pipeline. This function evaluates how the number of switching genes influences prediction accuracy and optionally visualises the results. It is typically used iteratively as part of a workflow to fine-tune model parameters.

The function takes a `SingleCellExperiment` object that has already been processed with the GeneSwitches package, where the data has been binarised using `binarize_exp()` and switching genes have been identified with `find_switch_logistic_fastglm()`. It then tests a range of values—selected by the user—and evaluates accuracy at each step.

During each iteration, the function takes the current value from `n_sg_range` and selects that number of the top switching genes. It then predicts cell positions and measures inaccuracies, storing the results in a data frame that maps the number of switching genes to their median inaccuracy.

If `plot = TRUE`, the function generates a plot showing how inaccuracy changes with the number of switching genes. The point with the lowest inaccuracy is labelled, this value is to be used when filtering the switching genes. If `plot = FALSE`, the function instead returns the data frame summarising the results.

This function helps optimise the selection of switching genes for PPR by identifying the number that minimises prediction error. It provides a straightforward way to assess accuracy across different gene selections.

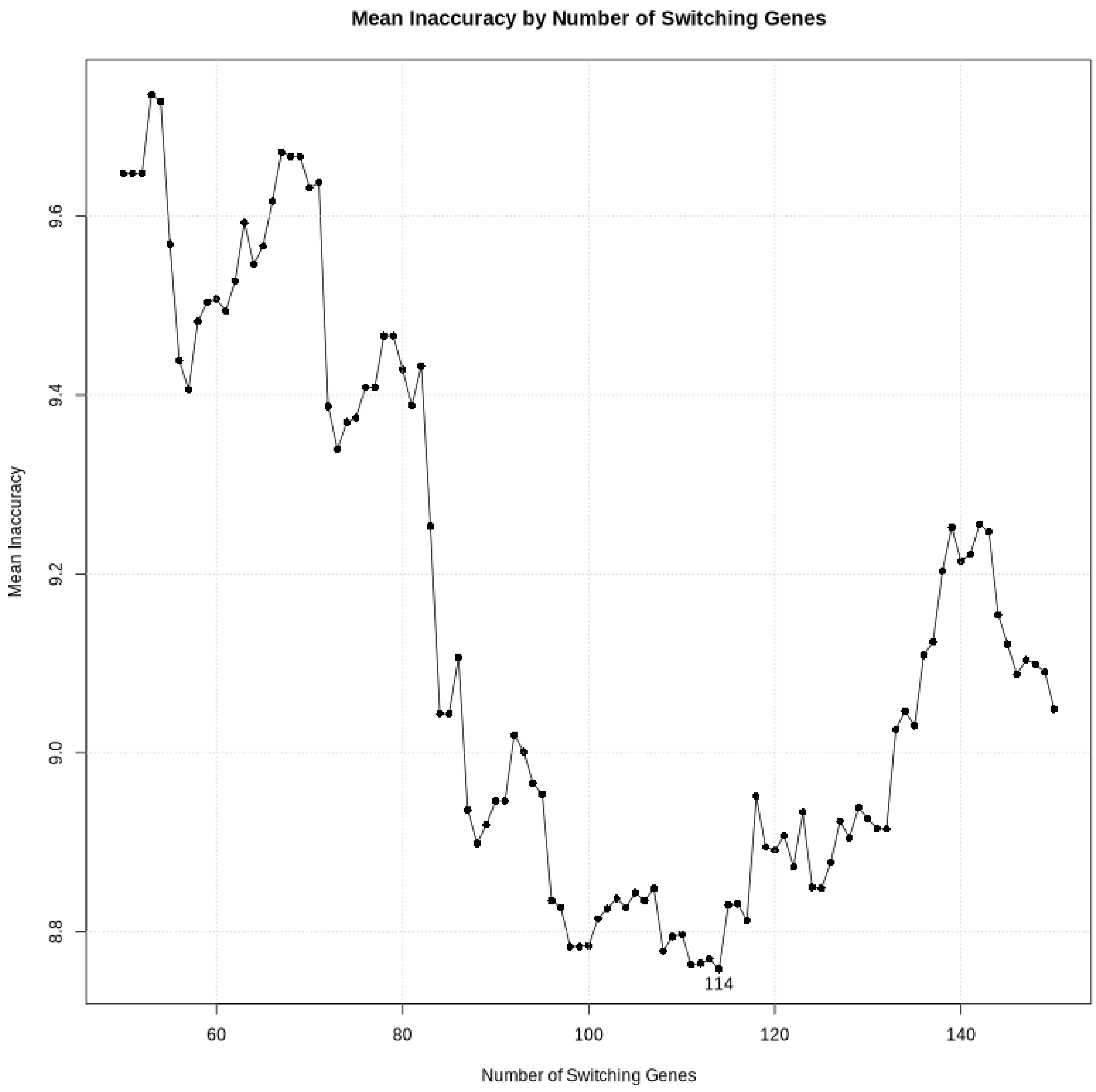


**Figure S3.** The effect of the number of switching genes on prediction accuracy in PPR. The `precision` function evaluates how varying the number of switching genes influences prediction error. The plot shows inaccuracy (y-axis) as a function of the number of switching genes (x-axis). The point corresponding to the lowest inaccuracy is highlighted (114 switching genes).

#### ppr_vioplot:

`Ppr_vioplot` generates a new plot,  rather than overlaying the `ppr_plot`. `ppr_vioplot` generates a violin plot to visualise predicted pseudotime positions for cells in a sample alongside reference cells, grouped by identity. The function facilitates comparison between the distributions of predicted cell positions (from the samples) and distributions of pseudotime values in the reference trajectory. It accepts a ppr object or a list of ppr_objects (samples_ppr), representing predicted pseudotime scores for cells from a sample. It also requires a SingleCellExperiment object (reference_sce) containing reference trajectory data. For data extraction, the function retrieves the highest PPR score positions (likely pseudotime positions) for each cell in samples_ppr. It then extracts true pseudotime values from the reference and maps them to indices along a 0–100 scale. Reference cells are grouped based on a selected identity. The plot is constructed using violin plots to represent pseudotime distributions. Reference groups (from reference_sce) appear in grey, while samples (from samples_ppr) are coloured using RColorBrewer palettes. The groups are ordered by mean pseudotime to ensure a logical layout.

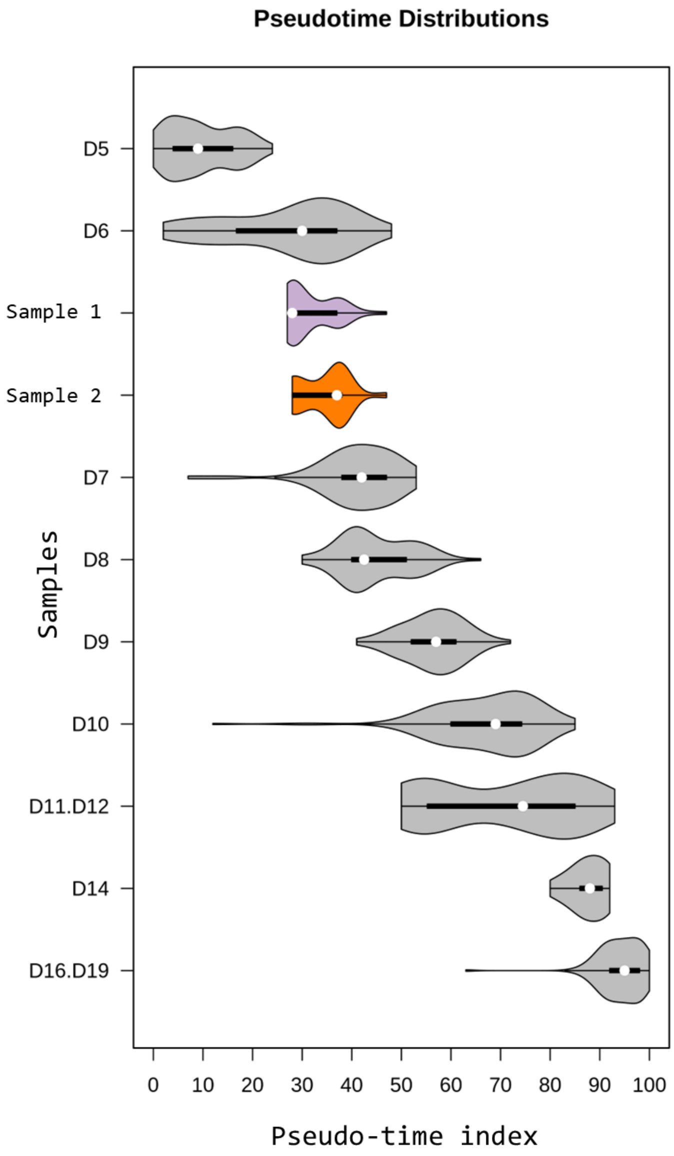


**Figure S4.** ppr_vioplot output plot.
Predicted position of two epiblast samples upon a development trajectory. Grey violin plots represent the true pseudotime of cells in the reference, split by a metadata value in this case day of development (D5, D6, ...). Purple and Orange violin plots represent the predicted pseudo-time for cells in
